# Arthropods on native versus alien woody plants: understanding variation across host plant taxonomy and geography

**DOI:** 10.1101/2025.09.04.674291

**Authors:** Colleen Whitener, Allen H. Hurlbert

## Abstract

Alien plants have generally been shown to support reduced arthropod abundance, biomass, and diversity, but inferences have typically come from studies limited in taxonomic and geographic scope. Here, we make use of data from a unique citizen science project, *Caterpillars Count!*, that consists of nearly 70,000 standardized surveys of woody plant branches across eastern North America on over 100 plant species. From these data we find that caterpillar (i.e. larval Lepidoptera) occurrence was nearly 4 times higher, beetle (order Coleoptera) occurrence was 50% higher, and spider (order Araneae) occurrence was 17% higher on native versus alien plants. The occurrence of hoppers and allies (suborder Auchenorrhyncha) was slightly higher on alien plants, while there was little difference in occurrence on native versus alien plants for ants (family Formicidae) or true bugs (suborder Heteroptera). Species richness of caterpillars, spiders, and beetles was also higher on average on native versus alien plants. The increased occurrence of caterpillars on native plants was consistent across plant families, whereas differences between native and alien plants for the other arthropod groups were highly variable across plant families. Caterpillar occurrence varied widely both within and between plant families, suggesting the importance of other plant species characteristics beyond native versus alien origin. Finally, we show that caterpillar occurrence appears to increase with latitude on alien plants at a faster rate than it does on native plants, but that this difference can be attributed to shifts in the identity and composition of plant species across the latitudinal gradient within our dataset. These findings have implications for how the increased spread of alien plants may impact ecosystems and food webs, and for land managers seeking to mitigate those changes.

## Introduction

One of the most pervasive human impacts on ecological systems is the introduction and facilitated spread of alien species (a term here used synonymously with ‘exotic’ or ‘non-native’; Simberloff et al. 2013). While some species like kudzu (*Pueraria montana*) and leafy spurge (*Euphorbia esula*) are well known for becoming widespread and causing significant economic damage (Pimentel et al. 2000), the vast majority of alien plants are not technically invasive and their individual impacts are often unknown. Nevertheless, the number of alien plant species becoming naturalized in North America has been increasing over time (Palmer 2005, van Kleunen et al. 2015, 2018), even in relatively pristine protected areas such as national parks (Miller et al. 2021). Collectively, it is clear that the rise of alien plant species has changed the diversity and composition of forests, savannahs, and grasslands, which may in turn have ecosystem-level effects including on soil characteristics (Diez et al. 2010, Duchicela et al. 2012), disturbance regimes (Mack and D’Antonio 1998), and even the facilitated colonization of other alien species (Jordan et al. 2008, Flory and Bauer 2014).

As primary producers and as the structurally dominant organisms in most ecosystems, woody plants additionally serve as both habitat and resources for a diverse array of consumer species. Forest arthropods in particular may be especially sensitive to changes in the identity of plant species in wooded environments. Many leaf chewers such as caterpillars (Lepidoptera larvae) have host plant associations such that they can only feed on plants of a particular species, genus, or family (Wagner 2005, Forister et al. 2015). Because these herbivores lack a shared coevolutionary history with alien plant species, they are often less able to tolerate or ameliorate the effects of the novel suite of secondary defensive compounds encountered in host tissue (Cappuccino and Arnason 2006). As a result, a number of studies have documented decreased abundance, diversity, and performance of these arthropod herbivores on alien species (Tallamy and Shropshire 2009, Burghardt et al. 2010, Tallamy et al. 2021, Narango and Straley 2025).

Although the effect of alien plants on arthropod abundance is expected to be greatest for herbivorous leaf chewers like caterpillars, other feeding guilds may be impacted as well (Lalk et al. 2021). Phloem and leaf suckers such as aphids (Hemiptera: Sternorrhyncha) and hoppers (Hemiptera: Auchenorrhyncha) exhibit some degree of host plant specialization in at least some systems (Cook and Denno 1994, Novotny et al. 2010). Ants have repeatedly evolved mutualistic associations with particular plant species, providing defense against herbivores in exchange for resources provided at extrafloral nectaries (Rico-Gray and Oliveira 2008, Chomicki and Renner 2015). Even some predators like spiders may have coevolved with native plants with respect to patterns of crypsis or the exploitation of particular plant architectures for web-building or hunting (Vasconcellos-Neto et al. 2017, Landsman et al. 2021, Hesselberg et al. 2023). However, tight obligate relationships may be more the exception than the rule in these latter groups. A meta-analysis found that vegetation plots that had been invaded by alien plant species did indeed have the strongest negative impacts on caterpillar abundance relative to uninvaded plots (67% of studies with decreases 0% with increased abundance; (Litt et al. 2014). In contrast, for beetles (Coleoptera), hemipterans, and spiders approximately half of studies showed decreased abundance in alien-invaded plots, while 25-33% of studies actually showed increased abundance (Litt et al. 2014). These results are consistent with a number of syntheses generally showing the strongest negative impacts for herbivores, but variable impacts on other ecological groups (Lalk et al. 2021).

The perceived effect of alien plants on arthropods can vary with the scale at which arthropod abundance or diversity is measured. Some studies like the ones summarized in the Litt et al. (2014) meta-analysis quantified arthropod abundance or diversity at the scale of vegetation plots or patches that may include multiple plant species (see also Carvalheiro et al. 2010, Narango et al. 2017). However, if the goal is to understand the specific effect a given plant species has on arthropod abundance locally, characterizing the arthropod response at the scale of a vegetation plot introduces other potentially confounding variables. For example, Landsman et al. (2021) found greater spider abundance in vegetation plots with an invasive grass than without, mostly due to the increased structural complexity provided by the alien plant cover that was not present in uninvaded plots. Other plot-level variables related to focal plant density or diversity may additionally impact the perceived arthropod response at this scale (Siemann et al. 1998, Haddad et al. 2009).

Another approach used in the literature for quantifying the ability of plant species to support arthropod diversity includes tabulating the total number of arthropod species known to associate with a given host species or genus across its range by synthesizing literature records (e.g., Tallamy and Shropshire 2009, Narango et al. 2020). While an exhaustive compilation such as this has the advantage of presenting the most complete picture possible of known host-arthropod associations across a broad suite of species, it may not provide an accurate characterization of the plant’s ability to support arthropods at the local scale. This is because the total number of arthropod associates known for a given host plant species is likely positively confounded with the geographic range size of the host (Leather 1986, Jones and Lawton 1991). For example, a widely distributed plant species and a narrowly distributed plant species might support the same average density and diversity of arthropods at a local scale, but the widely distributed species might have many more known arthropod associates simply because it encounters different arthropod communities in different parts of its range. In addition, a count of known host- arthropod associations will also be sensitive to the number and intensity of studies that have been conducted for a given host (e.g., Gregory 1990).

In contrast, studies that quantify arthropod response at the scale of individual plants or branches (e.g., Burghardt et al. 2010, Hartley et al. 2010, Parsons et al. 2020) most directly assess the contribution of plant species to local ecosystem function. Being able to infer the relative arthropod abundance and richness at this scale is not only critical for understanding the fundamental ecological and evolutionary processes governing biodiversity, but also for applications in ecological restoration where it is necessary to distinguish the plant species that might support more robust food webs in support of broader conservation goals (Tallamy 2007, Tallamy and Shropshire 2009). However, most such studies at the branch or individual scale typically examine just a small number of plant species, making it difficult to generalize more broadly about the effect of native and alien categories as a whole.

In addition, the vast majority of these studies have been conducted within a single site or region, which precludes the ability to evaluate whether any differences between native and alien species vary over large geographic gradients and are dependent on the climatic or regional context. Arthropod abundance and diversity are known to vary in general along gradients of elevation, temperature, and precipitation (Hysell et al. 1996, Progar and Schowalter 2002, Hodkinson 2005, Lessard et al. 2011, Ghosh-Harihar 2013). Latitudinal gradients encompass multiple sources of environmental variation, and the effect of alien host plant species could potentially vary latitudinally for two different reasons–(i) variation in climate across the latitudinal gradient could impact arthropod abundance and occurrence directly, independent of plant species, or (ii) the identity of plant species surveyed could vary across the latitudinal gradient, shifting toward species that host more or fewer arthropods on average.

Here, we present an analysis of foliage arthropod occurrence data collected as part of the citizen science project *Caterpillars Count!* (Hurlbert et al. 2019). The dataset includes information on the density and occurrence of caterpillars, beetles, spiders, true bugs, and hoppers for over 100 plant species sampled across a latitudinal gradient throughout eastern North America. Using this dataset which is unprecedented in geographic scope and taxonomic coverage, we ask the following questions. (1) To what extent do native plant species support higher arthropod occurrence and species richness compared to alien species, and how do these patterns vary across arthropod groups? (2) How do differences in arthropod occurrence between native and alien plant vary by plant family? For caterpillars in particular, we ask (3) how do individual plant species rank with respect to caterpillar occurrence, and what is the degree of phylogenetic signal in this trait across host plants?, and (4) Does the influence of host plant native status on the ability to support caterpillars vary latitudinally? We take advantage of this novel, geographically extensive dataset to address these questions.

## Methods

### Foliage surveys

We used foliage arthropod observations from the citizen science project *Caterpillars Count!* which enlists volunteers to monitor woody vegetation from across primarily eastern North America (Hurlbert et al. 2019). Participation revolves around monitoring sites, each of which has between 10 and 120 tagged survey branches. On a given survey date, all branches at the site are examined either using a beat sheet survey (striking a branch ten times with a stick over a white sheet; Montgomery et al. 2021) or a visual inspection of 50 leaves and associated twigs and petioles (as pioneered at Hubbard Brook Experimental Forest; Zammarelli et al. 2022). Each site chooses the survey method most appropriate for its aims, but arthropod density estimates have been found to be similar between methods (Hurlbert et al. 2019). The frequency of monitoring varies by site, but is typically weekly to monthly over the growing season. All arthropods observed are identified to order (or suborder within Hemiptera, family for ants), enumerated, and head-body length is estimated to the nearest mm. We focused on the following six arthropod groups that had sufficient observations for comparison: ants (Family Formicidae), beetles (order Coleoptera), caterpillars (order Lepidoptera), leafhoppers, treehoppers, and cicadas (hereafter “hoppers”, suborder Auchenorrhyncha), spiders (order Araneae), and true bugs (suborder Heteroptera).

To account for seasonality in arthropod abundance, we restricted analyses to surveys conducted during the months of June and July only, and excluded surveys whose leaves were indicated to be wet (e.g. from a recent rainstorm) which might have reduced arthropod occurrence. In total, we analyzed 68,741 branch surveys conducted on 363 different plant species from 212 sites in eastern North America (east of 100° W) ranging from 32.33° to 47.78° N latitude collected between 2010 and 2024.

We conducted within-family comparisons for the 10 plant families (Adoxaceae, Caprifoliaceae, Cornaceae, Malvaceae, Moraceae, Oleaceae, Rhamnaceae, Rosaceae, Sapindaceae, Ulmaceae) that had at least 60 surveys conducted on at least 7 distinct branches within each plant origin group. We calculated species-specific measures of caterpillar occurrence for the 100 host plant species that had at least 40 surveys conducted on at least 5 distinct branches. The median plant species in the dataset had 156 arthropod surveys conducted on 18 distinct branches.

### Native status

Plant native status was determined by the USDA PLANTS Database (Natural Resources Conservation Service 2024). Plant species were designated “native” to North America if their Native Status Code for North America or the Lower 48 States was listed as Native [“N”], Probably Native [“N?”], Native and Invasive [“NI”, meaning native in at least part of the region, but invasive in another], or Probably Native and Invasive [“NI?”]. Plant species were listed as “alien” if their Native Status Code for North America of the Lower 48 States was listed as Garden Persistent [“GP”, persists in gardens and around old habitations, not naturalized], Probably Garden Persistent [“GP?”], Invasive [“I”], Probably Invasive [“I?”], Waif [“W”, an ephemeral introduction, not persistently naturalized], or Probably Waif [“W?”].

### Plant phylogeny and phylogenetic signal

We used the ‘rtrees’ package in R (Li 2023) to create a phylogeny of our focal plant species based on the megatree for vascular plants from Jin and Qian (2022). As a measure of phylogenetic signal in caterpillar occurrence across host plants, we calculated Blomberg’s K (Blomberg et al. 2003) and compared it with a null distribution generated from 999 randomized permutations of trait values across the tree tips.

### Arthropod identification and finer taxonomic identification

As part of each foliage survey, users could optionally submit photos of the arthropods they observed. Photos shared as a part of *Caterpillars Count!* surveys get automatically shared to the project iNaturalist account where the community of identifiers (Campbell et al. 2023) could potentially 1) confirm or reject the order-level classification of *Caterpillars Count!* users, and 2) provide finer resolution taxonomic identifications. Of all arthropod photos for which an order-level classification could be made, *Caterpillars Count!* users were correct in their classification of these six arthropod groups 93% of the time (unpublished data).

We also tallied the number of unique taxa observed on each host plant species as a function of the total number of surveys for which at least one arthropod photo was included. *Caterpillars Count!* participants are strongly encouraged to submit photos of caterpillars, and thus this taxonomic group has the greatest potential for species identifications, but each of the six arthropod taxa had at least 1,000 specimen photos. The number of unique taxa could include observations that were only identified to the genus or family level, as long as there were no other identified taxa counted that were also within that genus or family.

### Statistical analysis

For each plant species, plant family, and native status group, we calculated the proportion ( ) of branch surveys which recorded at least one individual of the focal arthropod group of interest. We focus on the proportion of surveys, i.e. occurrence, rather than density per survey because of the right-skewed distribution of abundance values and to reduce the impact of those rare, high value outliers. The margin of error for the sample proportion was calculated as 1.96 * (( * (1 -) / *n*)^0.5 where *n* is the number of surveys, allowing the calculation of 95% confidence intervals. We conducted a two-sample proportion test to compare the proportion of surveys on native versus alien plants for each arthropod group and within each plant family. We conducted such comparisons as long as there were at least 10 surveys per group. This meant no comparison between alien and native species was conducted for some families like Fagaceae because there were insufficient surveys on alien species within the dataset.

For evaluating latitude as a covariate, we compared two hierarchical mixed effects models of the binomial response of caterpillar presence with latitude, native plant status, and their interaction as predictors. Both models accounted for the fact that individual survey branches within a monitoring site will generally share the same unmeasured climate and landscape characteristics that could potentially influence arthropod occurrence (i.e., site was included as a random effect), and one of the models additionally accounted for plant species as a random effect. We conducted both models because most plant species occupy only a subset of the full latitudinal gradient. The first model evaluates the difference in caterpillar occurrence between native and alien species across the latitudinal gradient given the existing plant species occurrences, whereas the second model evaluates whether there is a consistent effect of latitude on caterpillar occurrence when controlling for intercept differences between individual plant species. Latitude values were centered and scaled to minimize errors with model convergence.

## Results

### Overall arthropod occurrence and richness

The effect of plant species native status on arthropod occurrence varied by arthropod group (Figure 1). The greatest effect was for caterpillars, which were found on 8.1% of surveys on native plant species, but only 2.2% of surveys on alien plants (*p* < 10^-10^). Beetles were found on 21.9% of native surveys, but only 14.7% of alien surveys (*p* < 10^-10^), and spiders were found on 20.0% of native surveys and 17.1% of alien surveys (*p* < 10^-4^). The other 3 arthropod groups had very similar estimates of occurrence on native and alien surveys, with some evidence that hoppers were present on a slightly greater proportion of alien plant species branches (13.7% versus 12.2%, *p* = 0.014).

**Figure 1.**
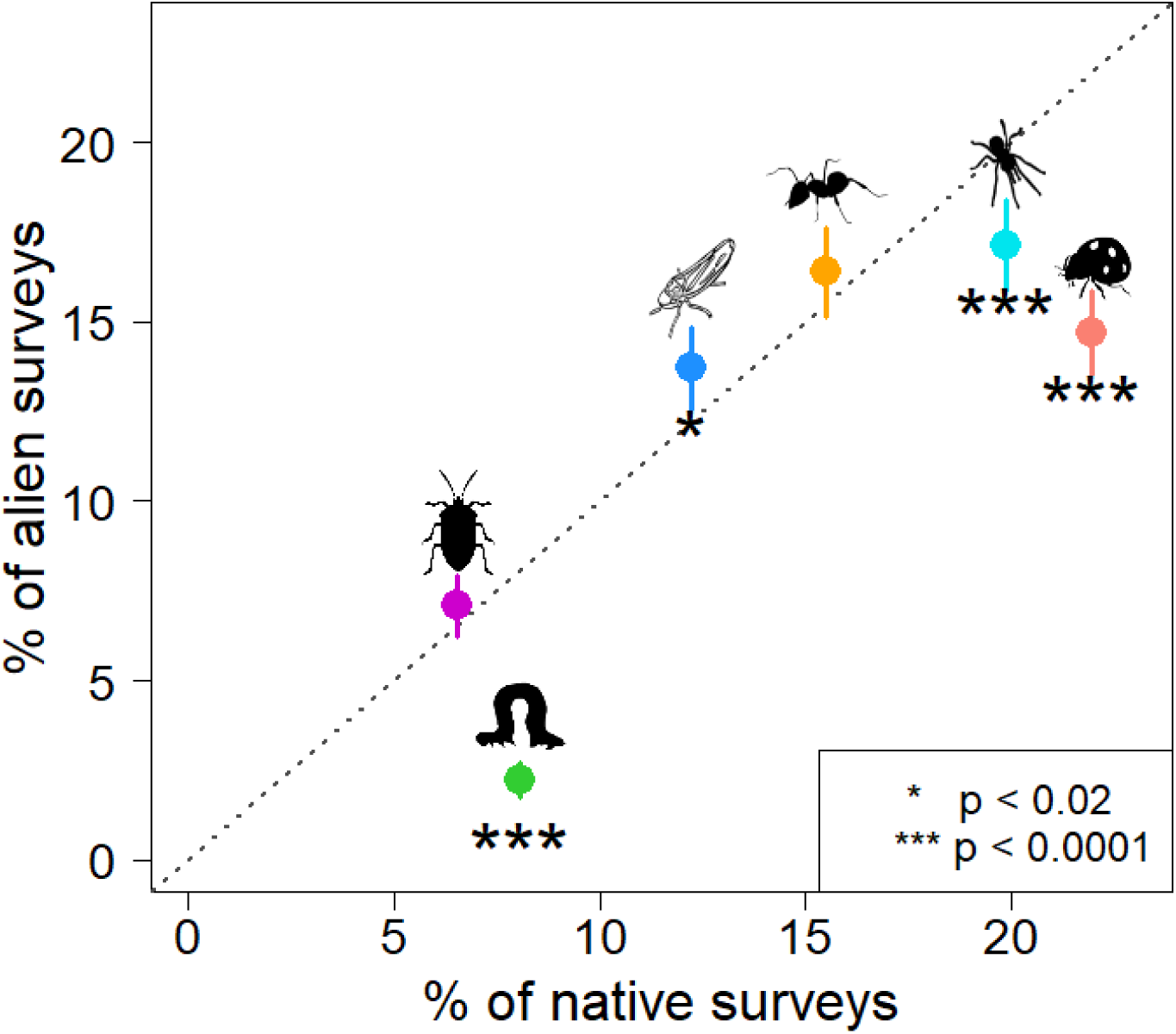
The proportion of surveys on which different arthropod groups were observed on either native (*n* = 64,964 surveys) or alien (*n* = 3,428 surveys) plant species. Line segments indicate the 95% confidence interval. Confidence intervals on the % of native surveys are typically smaller than the plotting symbol due to the high sample size. Dotted line displays y = x for reference. Illustration credit: A. Hurlbert.

In addition to having higher caterpillar occurrence on average, native plant species also supported a higher diversity of caterpillar species, even controlling for the fact that surveys on native plants tend to have more surveys with photos than surveys on alien plants (Figure 2).

**Figure 2.**
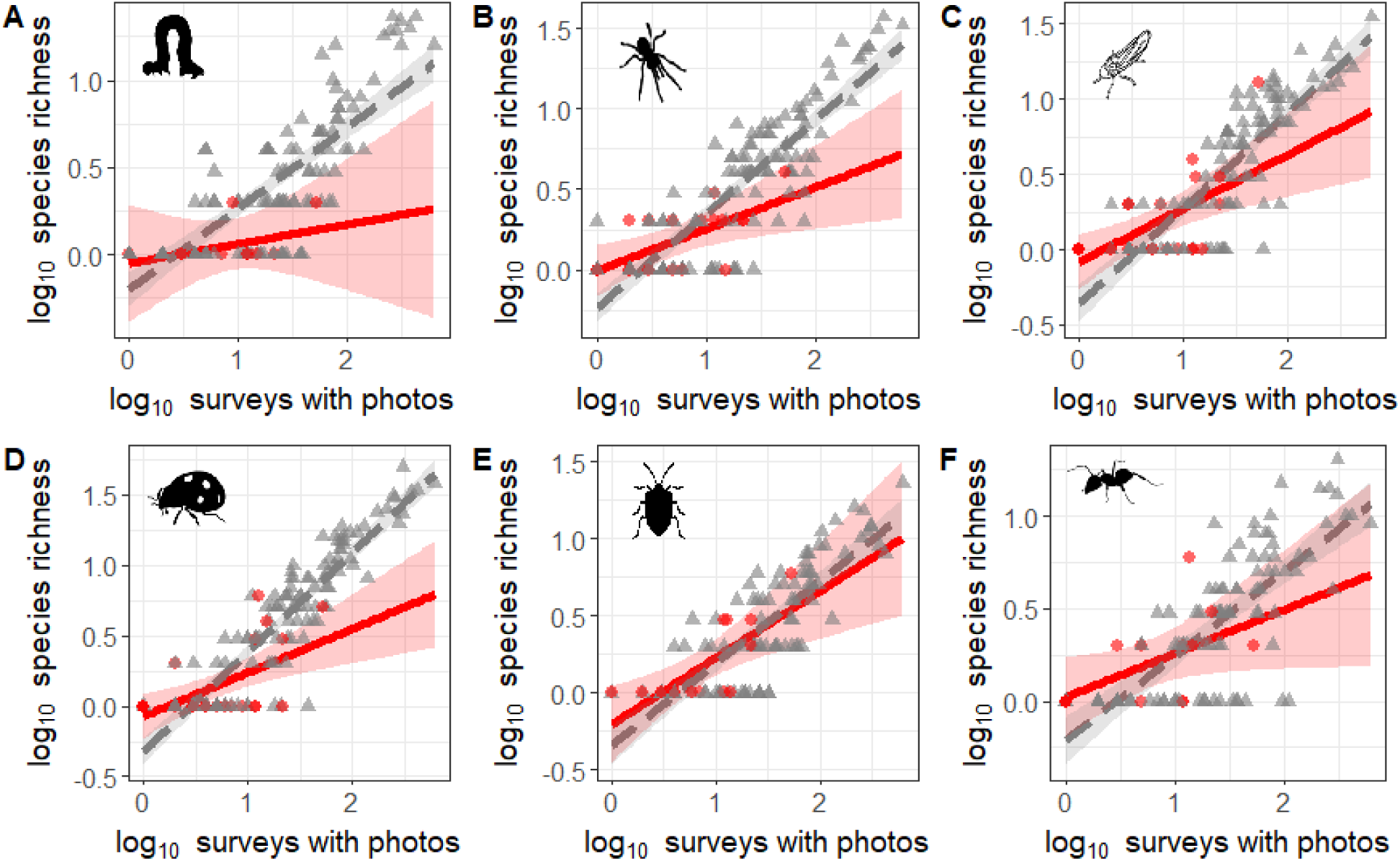
Number of unique taxa (or unique genera, subfamilies, or families) of (a) caterpillars, (b) spiders, (c) hoppers, (d) beetles, (e) true bugs, and (f) ants observed on a given host plant species as a function of the number of surveys with arthropod photos for that plant species. The number of species increases more steeply for native (gray) than alien (red) plant species for all groups except true bugs (see text). Illustration credit: A. Hurlbert.

Specifically, the number of caterpillar species increased more steeply for native than alien plant species as the number of surveys increased (slope difference = 0.35, *p* = 0.038). On average, given 100 surveys for which the user documented arthropods with photos, one would expect only 1.5 caterpillar species on an alien host plant, but 5.4 species on a native host plant.

Strong patterns of higher species richness on native versus alien plants were also observed for spiders ( = 0.33, *p* = 0.001) and beetles ( = 0.40, *p* < 0.0001), slightly weaker patterns were observed for hoppers ( = 0.28, *p* = 0.015), and and no strong difference was observed for true bugs (*p* = 0.46) or ants (*p* = 0.08).

### Arthropod occurrence by plant family

Differences in the occurrence of arthropods between native and alien plant species were more apparent within certain plant families compared to others (Figure 3). The most uniform positive effect of native status was for caterpillars, where six of the ten families compared exhibited evidence of greater caterpillar occurrence on native species. The greatest effects were for Sapindaceae (maples, buckeyes; *p* < 10^-45^), Cornaceae (dogwoods; *p* < 10^-17^), Oleaceae (ashes, privets, lilacs; *p* < 10^-9^), and Malvaceae (mallows, lindens; *p* < 10^-5^), for the most part families with the largest sample sizes.

**Figure 3.**
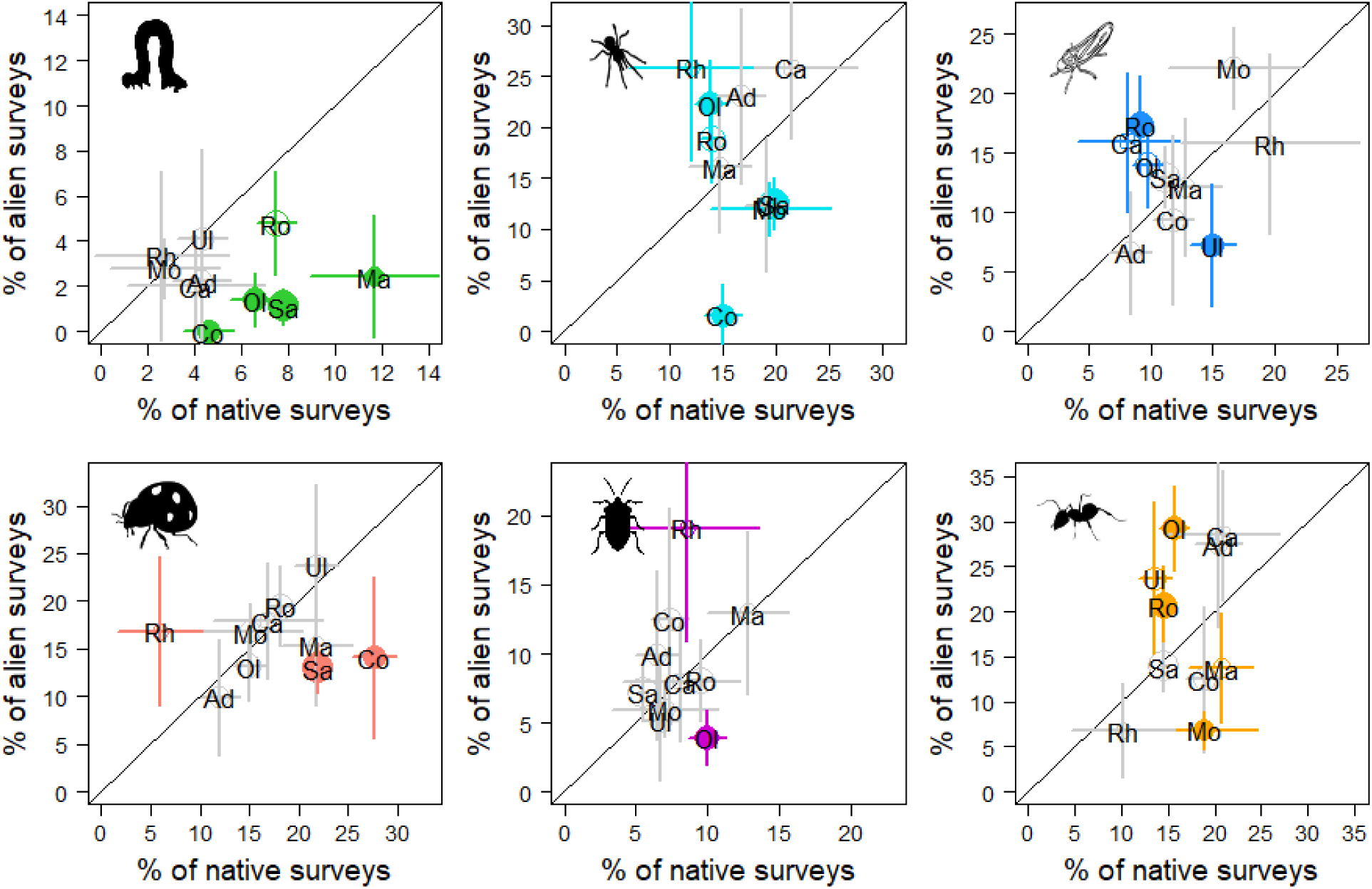
Comparisons of arthropod occurrence (mean and 95% CI) on native versus alien plant species within each of ten plant families for (a) caterpillars, (b) spiders, (c) hoppers, (d) beetles, (e) true bugs, and (f) ants. Comparisons for which the two-sampled proportion test *p* < 0.01 are indicated with solid colored circles, and symbol size reflects the branch sample size. Plant family abbreviations, along with the number of native and alien survey branches, respectively, are as follows: Ad = Adoxaceae (1024, 91), Ca = Caprifoliaceae (172, 151), Co = Cornaceae (1545, 64), Ma = Malvaceae (531, 124), Mo = Moraceae (185, 574), Ol = Oleaceae (2110, 359), Rh = Rhamnaceae (117, 89), Ro = Rosaceae (3778, 334), Sa = Sapindaceae (14,150, 641), Ul = Ulmaceae (1409, 97). Illustration credit: A. Hurlbert.

For the remaining arthropod groups, there was mixed evidence for the effect of native status across plant families. Beetles and spiders both had higher occurrence on native members of Sapindaceae (*p* < 10^-7^) and Cornaceae (*p* < 10^-2^), but lower occurrence on native members of at least one other family. The occurrence of true bugs was very similar between native and alien members of most families with the exception of Oleaceae (native > alien; *p* < 10^-6^) and Rhamnaceae (buckthorns, alien > native; *p* = 0.03). The greatest effect of native status for hoppers was for the Rosaceae (plums, cherries, crabapples, roses; alien > native; *p* = 0.0001).

Ant occurrence was much higher on alien members of Oleaceae (*p* < 10^-7^) but native members of Moraceae (mulberries; *p* < 10^-4^).

Just as the effect of native status was mixed across families for any given arthropod group, the effect of native status was also often mixed across arthropod groups within some plant families. For example, native members of Oleaceae supported more caterpillars (*p* < 10^-9^) and true bugs (*p* < 10^-6^), but fewer spiders (*p* < 10^-3^) and ants (*p* < 10^-7^) than alien species. In contrast, for both Sapindaceae and Oleaceae, the only differences observed for which *p* < 0.01 all indicated greater arthropod occurrence (caterpillars, beetles, and spiders) on native species, with no arthropod groups having substantially lower occurrence on native species.

### Caterpillar occurrence on individual host plant species

Although there were clear signals of plant native status on caterpillar occurrence overall, as well as within plant families, one obvious source of variation was between individual species within either alien or native plant origin groups (Figure 4; Appendix S1: <u>Table S1</u>). Amongst native plant species, caterpillar occurrence varied from 0% (*Malus angustifolia, Carya tomentosa*, and *Aesculus parviflora*) to 19.4% (*Hamamelis virginiana*). For 14 of the 15 alien species with sufficient surveys (Figure 4, in red), caterpillars were found on no more than 3.8% of survey branches. The alien species with the highest caterpillar occurrence, *Ulmus pumila* (6.4%), ranked below 55 native species in our dataset.

**Figure 4.**
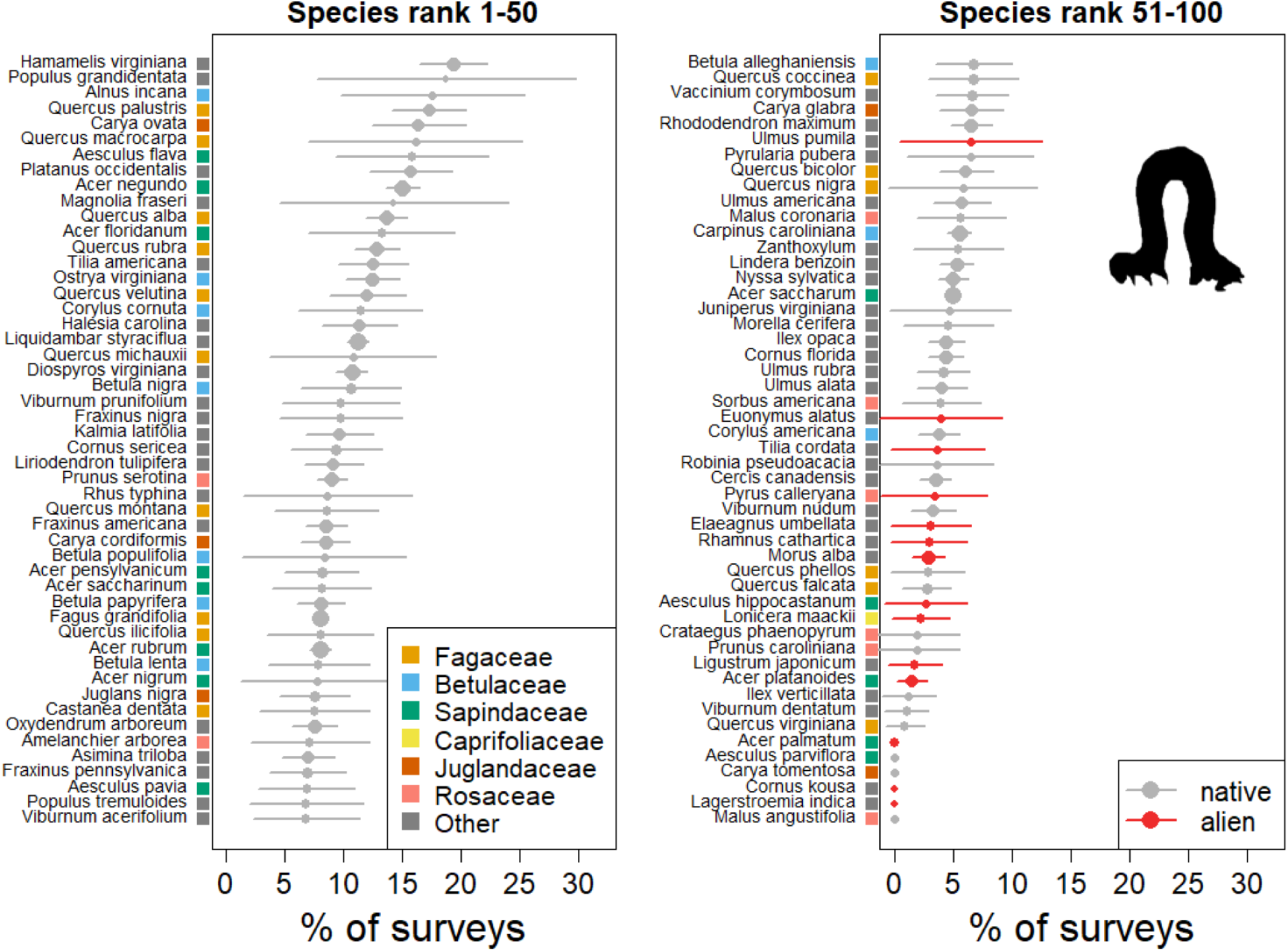
The percentage of foliage surveys on which a caterpillar was observed (plus 95% CI) for the 100 plant species with at least 40 surveys. Native species are plotted in gray, and alien species in red. Species names are color coded according to the 6 most common plant families. Symbol size is scaled to reflect sample size of branch surveys. Illustration credit: A. Hurlbert.

Caterpillar occurrence exhibited very little phylogenetic signal (Blomberg’s *K* = 0.04, *p* = 0.77). The top seven plant species represent six different plant families, and nearly every plant family includes species in both the top 20% and bottom 20% of the rankings (Figure 4).

### Caterpillar occurrence over a latitudinal gradient

Finally, we examined how caterpillar occurrence differs between native and alien plant species across a latitudinal gradient. Ignoring variation due to plant species, we find that while caterpillar occurrence is higher for native than alien plants on average (*p* = 1.3e-13), there is an indication that the way occurrence varies with latitude differs between native and alien host plant status (Figure 5, native status x latitude interaction term, *p* = 0.06). Specifically, caterpillar occurrence increases more steeply with latitude on alien plants than native plants, meaning that the difference between native and alien plants is smaller at higher latitudes.

**Figure 5.**
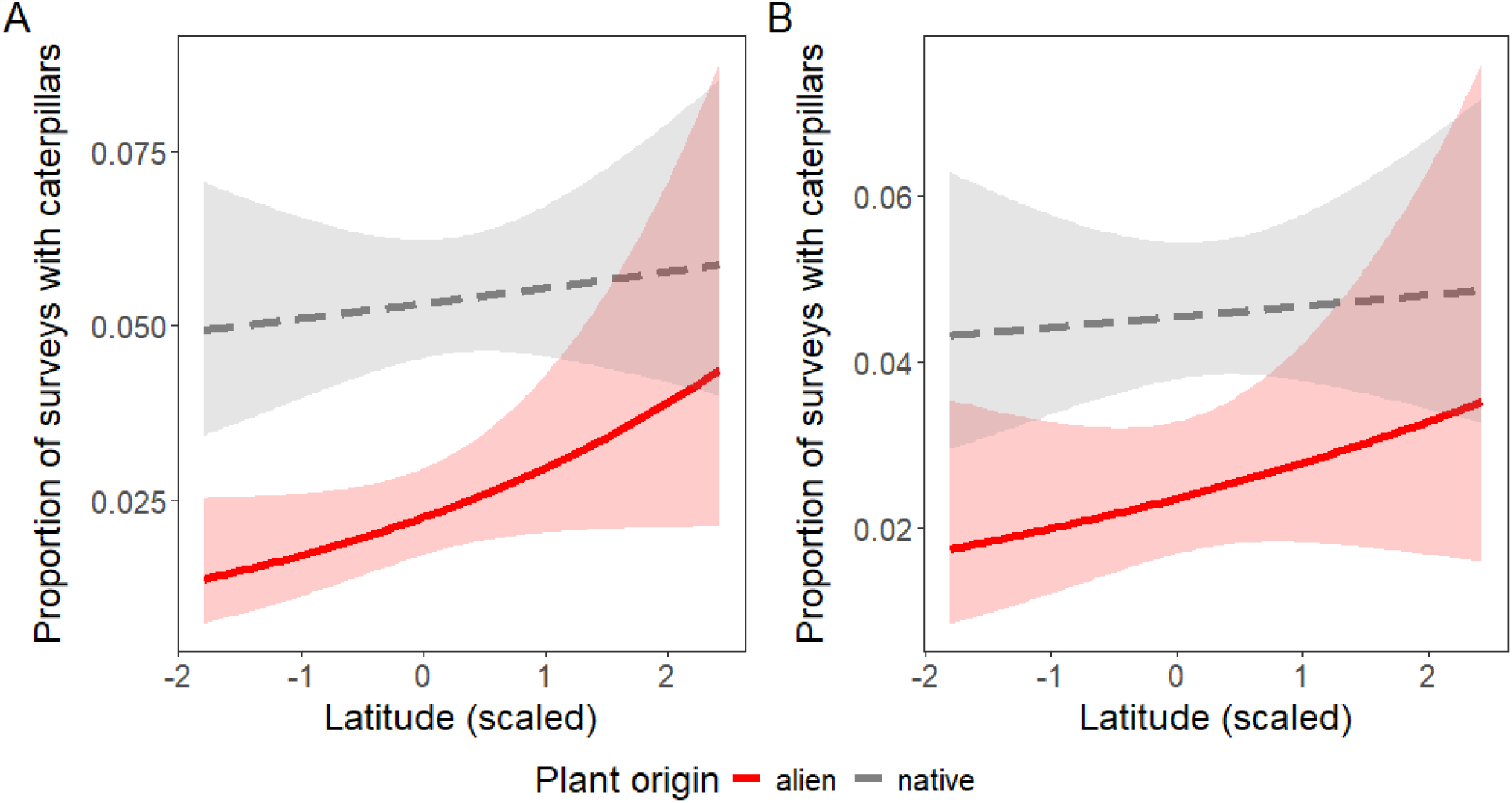
(a) Modeling the interaction between latitude and host plant origin when including site as a random effect reveals caterpillar occurrence increases with latitude, and is higher on native plants (*p* < 10^-11^), but the rate at which caterpillar occurrence increases with latitude is different on native (gray) versus alien (red) host plants (*p* = 0.02), with a greater effect of native plant origin at low latitudes. (b) A similar model that additionally included plant species as a random effect found a strong effect of host plant origin overall (*p* = 0.006) but no evidence of an interaction between latitude and host plant origin (*p* = 0.22).

When including plant species as a random effect in the model, there was still a strong effect of native plant status on average (*p* = 2.9e-5), however, uncertainty in the latitude and latitude x host plant origin status terms increased (*p* > 0.3 for both; Figure 5b). Thus, there was no consistent effect of latitude on caterpillar occurrence *within* plant species, regardless of native plant status. We show this directly for 8 geographically widespread plant species in Appendix S2: Figure S1. The effect of latitude on caterpillar occurrence is therefore due at least in part to compositional shifts in vegetation sampled across the latitudinal gradient.

## Discussion

Using a large-scale citizen science dataset on foliage arthropods, we showed how the extent to which native plants support more arthropods than alien plants varies by arthropod group, host plant taxonomy, and latitude. As expected, the effect of host plant origin was strongest for caterpillars, which were nearly 4 times more common, and which had the greatest difference in diversity, between native and alien plants. This effect of plant origin on caterpillars was greater than the 70-90% increase documented by Narango et al. (2017) and Parsons et al. (2020), but in line with the 3-4-fold increase in caterpillar density on native plants documented by Burghardt and colleagues (2010). Beetles, the majority of which are also phytophagous, had the next largest positive effect of native plant status on both occurrence and richness, followed by spiders with 19% higher occurrence on native plants. In contrast, ants, true bugs, and hoppers all had very similar levels of occurrence on native and alien plants, although ants and hoppers did have slightly higher diversity on native plants. While these differences between groups are partly explained by trophic guild, the stronger response of predatory spiders compared to phloem and leaf sucking hoppers was unexpected.

These overall effects of plant origin on arthropod occurrence are informative as they reflect the prevalence of the actual native and alien species present on the landscape of these monitoring sites. However, family-level variation in plant traits that influence host plant quality (e.g. chemical defenses, leaf nitrogen, physical structure) could potentially confound the interpretation of plant origin effects given that alien plants tend to be a non-random subset of the native flora phylogenetically (Diez et al. 2008, Cadotte et al. 2009). Comparing arthropod occurrence on native and alien species within plant families potentially controls for some of these sources of variation, and reveals that while the effect on caterpillar occurrence is fairly consistent across plant families, the effect on other arthropod groups is family specific. For example, the increased occurrence of both beetles and spiders on native plants was driven primarily by host plants in the Cornaceae and Sapindaceae families, whereas occurrence was actually higher on alien plants within Rhamnaceae, and in the case of spiders, Oleaceae.

The family-specific nature of many of these effects suggests that additional factors beyond host plant origin undoubtedly play a role. Some of the families in our dataset are primarily represented by just one or two species (e.g. 95% of surveys of alien Cornaceae were on either *Cornus kousa* or *Cornus mas*, 96% of surveys on alien Rhamnaceae were on *Rhamnus cathartica*, Appendix S1: Table S1), and so some of these family effects may in fact be due to the characteristics of individual plant species rather than family-level characteristics. We examined species level variation in detail for caterpillar occurrence, and found substantial variation within plant families in the ability of a species to support caterpillars, and very little phylogenetic signal. For example, native members of the genus *Quercus*, which is generally considered to support the highest diversity of Lepidoptera in North America (Narango et al. 2020), ranged uniformly in ranking from 4th to 97th out of the 104 species examined. More research is needed to explore the various functional (e.g., resource quality and defense, Mattson 1980, Carmona et al. 2011), macroecological (e.g., abundance, range size, dilution effects; Strong 1974, Leather 1986, Gougherty and Davies 2024), and macroevolutionary (e.g., clade age or size, phylogenetic distinctness; Connor et al. 1980, Novotny et al. 2010, Ness et al. 2011) traits that might explain this interspecific variation.

One of the advantages of this large citizen science dataset is the opportunity to evaluate broad geographic patterns in arthropod occurrence based on a standardized sampling methodology. In this case, we found that the difference between native and alien plants in their ability to support caterpillars varied with latitude. Generally, caterpillar occurrence increased slightly with latitude from Georgia (∼32° N) to Ontario (∼46° N), with the difference between native and alien plants greatest at low latitudes. However, taking into account the identity of individual host plant species which do not all occur uniformly across the latitudinal gradient, there was neither a strong latitude effect nor latitude by host plant origin interaction. This reinforces preliminary findings (Di Cecco and Hurlbert 2022) that patterns of caterpillar occurrence versus latitude are idiosyncratic within individual host plant species. Nevertheless, land managers in more southern regions should be aware of the particular benefit of replacing alien with native plants for supporting arthropod biodiversity, assuming that the alien species in our dataset are representative of what is commonly found on the landscape.

Although data from the *Caterpillars Count!* citizen science project are collected primarily by amateur naturalists and do not usually provide species-level identifications (only ∼3% of arthropod observations), they are a unique source of information for better understanding arthropod abundance, occurrence, and biomass in forest ecosystems. As the dataset continues to grow, both in geographic coverage and in the duration of long-term time series, we expect it to yield further insights into not just the importance of native versus alien species, but also how arthropod abundance and phenology are changing over time, and how they vary across finer-scale gradients in land use and urbanization. Ultimately, standardized large-scale surveys for arthropods will greatly increase our ability to understand and mitigate the various anthropogenic and environmental drivers of insect decline (Wagner et al. 2021).

## Acknowledgments

We would like to acknowledge the thousands of *Caterpillars Count!* participants who helped collect the data that made this research possible. This research utilized infrastructure developed through previous support from the National Science Foundation under award EF 1702708. Nosa Osawe provided helpful feedback on an earlier draft.

## Author Contributions

Allen H. Hurlbert obtained funding, conceived the study, and maintained infrastructure supporting project data collection. Colleen Whitener contributed to data collection, wrote the first draft, and conducted initial analyses. Allen H. Hurlbert revised analyses and wrote the second draft. All authors read and approved the final manuscript.

## Appendix S1

Table S1. The 100 plant species included in the species-level comparison, their native or alien status, prevalence in the dataset by number of surveys, branches, and monitoring sites, as well as the percent of surveys on which a caterpillar was observed, along with the 95% confidence limits on that percent.

Table S1 is available for review in the Github repository here: https://github.com/hurlbertlab/caterpillars-on-plants/blob/main/data/Table_S1.csv

## Appendix S2

Open Research Statement

All data and code underlying the analyses are available at http://github.com/hurlbertlab/caterpillars-on-plants [to be archived on Zenodo upon acceptance].

**Figure S1.**
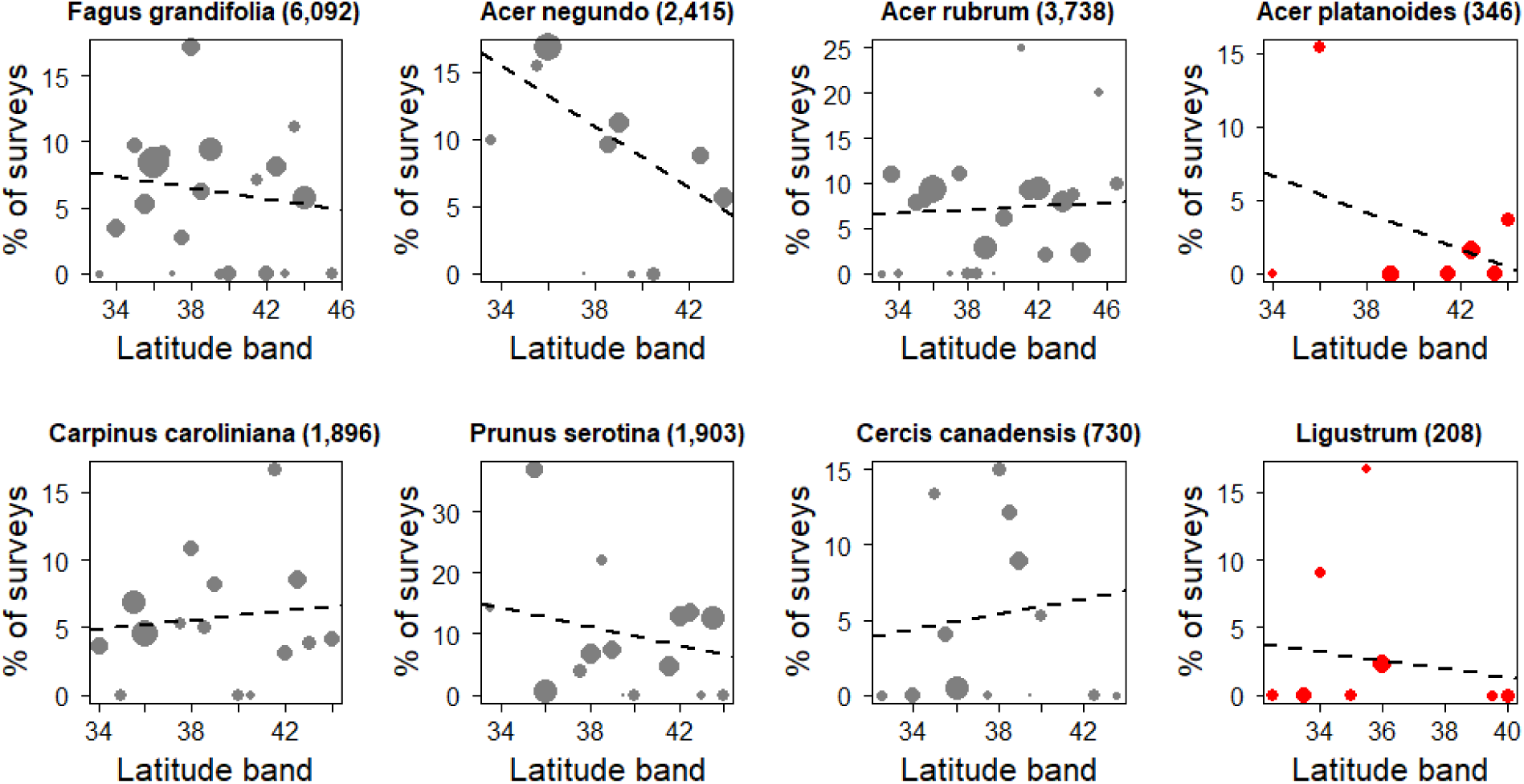
Across six native (gray) and two alien (red) host plant species, there is no consistent trend between caterpillar occurrence and latitude (*p* > 0.05 for all species).

